# Mapping of residues in leishmanial glyceraldehyde-3-phosphate dehydrogenase crucial for binding with 3’-UTR of TNF-α mRNA

**DOI:** 10.1101/2025.05.02.651801

**Authors:** Puja Panja, Sumit Das, Yuthika Dholey, Gaurab Chowdhury, Subrata Adak

## Abstract

Recently, we described that glyceraldehyde-3-phosphate dehydrogenase from *Leishmania major* (LmGAPDH) was present in extracellular vesicles and it inhibited host TNF-α expression during infection via post-transcriptional repression. The LmGAPDH binding with the AU-rich elements in 3’-untranslated region of TNF-α mRNA (TNF-α ARE) is sufficient for limiting this cytokine production, but the TNF-α ARE binding residues in LmGAPDH are still unexplored. RNA electrophoretic mobility shift assay (REMSA) and catalytic activity measurement revealed that the inhibition by TNF-α ARE was competitive with respect to cofactor NAD^+^ in LmGAPDH. To identify the TNF-α ARE binding residues of the LmGAPDH, we exploited a systematic mutational analysis of its NAD^+^ binding domain. Catalytic activity measurement indicates that both R13 and N336 amino acids in the NAD^+^ binding site are absolutely required for activity whereas other mutants including I14A, R16A, D39A and T112A showed higher Km (lower affinity) value for NAD^+^ binding and lower catalytic activity. REMSA studies revealed that the replacement of Arg-13 with Ala/Lys or Asn-336 with Ala resulted in complete loss of binding with the TNF-α ARE. I14A, R16A, D39A and T112A residues at or near NAD^+^ binding site showed lower binding with the TNF-α ARE compared to the wild-type protein. The protein induced fluorescence enhancement (PIFE) studies and *in vitro* protein translation assay further confirmed the REMSA results. Based on our findings, the NAD^+^ binding residues in LmGAPDH are important for TNFα ARE binding.

## Introduction

One of the classical glycolytic enzymes is glyceraldehyde-3-phosphate dehydrogenase (GAPDH), which catalyzes the reversible oxidative phosphorylation of glyceraldehyde-3-phosphate to the higher energy intermediate 1,3-bisphosphoglycerate in the presence of inorganic phosphate and nicotinamide adenine dinucleotide (NAD^+^). However, emerging evidences suggest that GAPDH is a moonlighting protein, which exhibits multiple non-glycolytic functions in different subcellular organelles [1, 2]. Apart from glycolytic function, membrane-bound GAPDH displays an important function for membrane fusion, endocytosis and iron transport [3–5]. Cytosolic GAPDH controls the stability of mRNA, post-transcriptional regulation and translation [6–10], and is required for heme insertion of iNOS [11] and ER to Golgi trafficking [12]. GAPDH in nucleus is involved in transcriptional regulation of gene, the preservation of DNA integrity, cell death through apoptosis, as well as nuclear tRNA export [13–17]. Unexpectedly, numerous human diseases, like tumorigenesis, diabetes and neurodegenerative disorders are dependent on GAPDH structure and function [18, 19].

Recently, we have shown that the GAPDH protein from *Leishmania major* (LmGAPDH) is localized within the extracellular vesicles (EVs) [20]. *Leishmania* EVs are mainly involved in the transporting of various biomolecules into host macrophage during infection [21–23]. By comparative studies among LmGAPDH overexpression, half knockout (HKO), and complement *Leishmania* promastigotes, HKO cells exhibited lesser virulence property compared to other cell lines when BALB/c mice were inoculated separately with all kinds of cell lines [20]. In addition, ELISA, RT-PCR, and immunoblot data showed that higher TNF-α protein expression occurred during infection of host macrophages with HKO cell lines and its EVs. *In vitro* protein translation studies suggested that the repression of TNF-α expression is directly proportional to LmGAPDH concentration. Furthermore, RNA electrophoretic mobility shift assay (REMSA) studies suggested that LmGAPDH has ability to bind with the AU-rich 3’ -UTR region of TNF-α mRNA (TNF-α ARE) [20].

GAPDH lacks canonical RNA-binding consensus sequences, for example RNP motifs, RGG boxes, KH domain or any other known nucleic-acid-binding motifs [24]. It is well-known that AU-rich sequence elements of mRNA are involved in GAPDH binding [25]. The inhibition study of the RNA-GAPDH complex formation by NAD^+^, NADH and ATP further indicated that the NAD^+^-binding site or (di) nucleotide-binding fold (Rossman fold) is involved in RNA binding [20, 25]. In addition, GAPDH has ability to interact with other RNAs including tRNA, rRNA and viral RNA [16, 26, 27]. A group of researchers established by using deletion mutant that the N-terminal 43 amino acid residues (Rossman fold) of human GAPDH (with GST fusion protein) are sufficient for TNF-α ARE binding [28] but the precise binding residue(s) in GAPDH is/are still unexplored. Another group of researchers suggested that the TNF-α ARE-binding site in GAPDH would span beyond the Rossman fold area including the dimer and tetramer interface [29]. Although structure of NAD^+^ bound GAPDH are now available [30–34], a high-resolution X-ray structure of RNA bound GAPDH is still unknown in the literature. Here we have identified TNF-α ARE binding residues in LmGAPDH by performing a logical mutational analysis in the NAD^+^ binding domain of this enzyme.

## Results

### Interaction of LmGAPDH and TNF-α ARE determined by RNA electrophoretic mobility shift assay (REMSA)

To determine apparent dissociation constant (K_D_) of wild type LmGAPDH protein for TNF-α ARE, we have performed the LmGAPDH dependent REMSA study. Our REMSA studies revealed that the binding of LmGAPDH with TNF-α ARE occurs in concentration dependent manner (Figure 1A), which is comparable to earlier studies [20, 25, 29]. Here we measured K_D_ of wild type LmGAPDH protein for TNF-α ARE by plotting % shifted band intensity of protein-RNA complexes versus the concentration of LmGAPDH. The K_D_ value of wild type LmGAPDH protein for TNF-α ARE is 1.7 ± 0.05 µM, which is comparable to previous studies [29]. Figure1B showed that increasing concentrations of cofactor NAD^+^ decreased the formation of LmGAPDH -TNF-α ARE complex. The Ki value of NAD^+^ against protein-RNA complexes formation is 0.48 ± 0.15 mM. This result suggests that the nature of the interaction of LmGAPDH with the TNF-α ARE might be reversible as well as of competitive nature.

**Figure 1.**
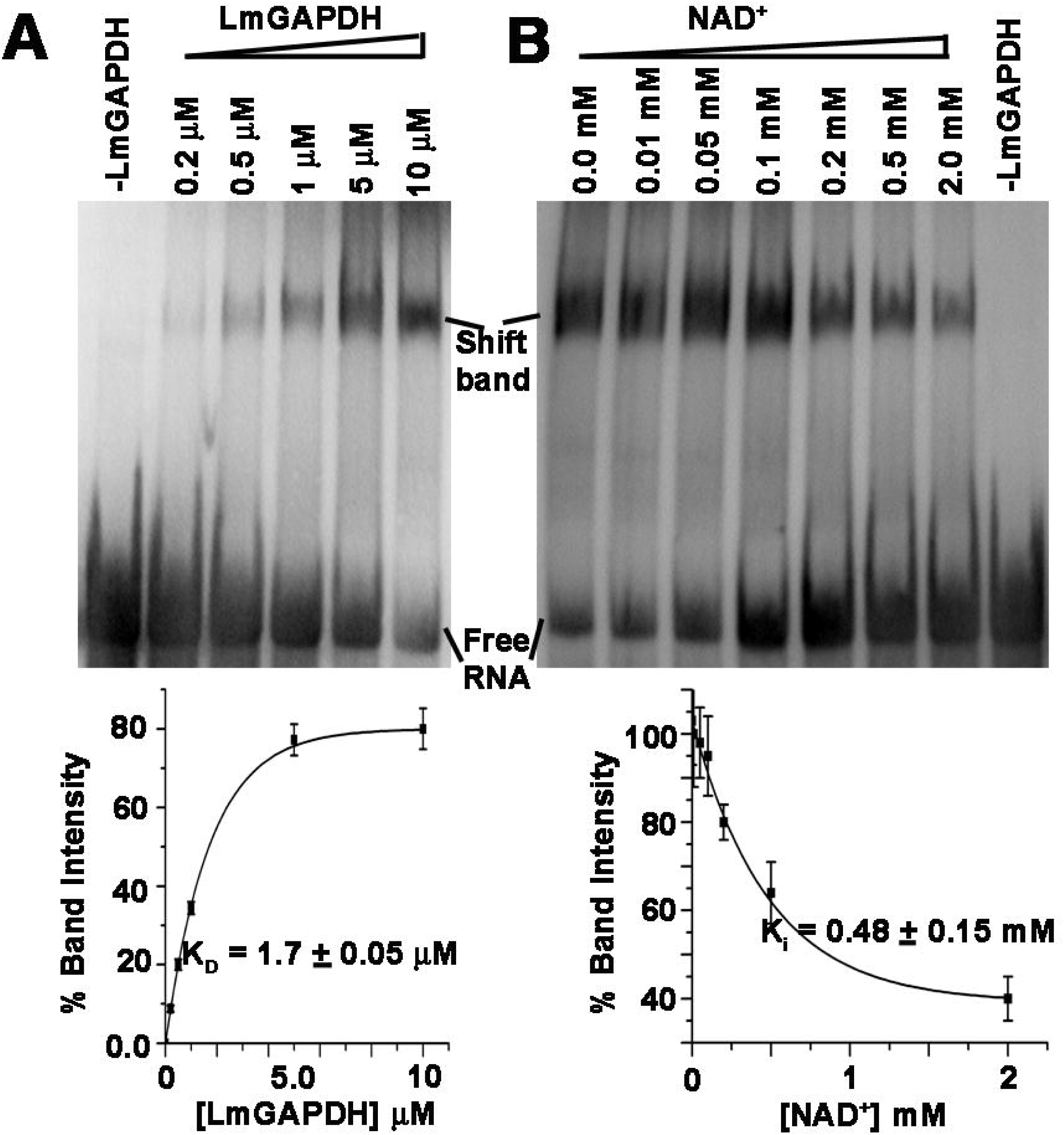
LmGAPDH dependent TNF-α ARE-enzyme complexes in presence and absence of NAD^+^. Panel A (Upper panel), denoted the RNA electrophoretic mobility shift assay (REMSA) image for LmGAPDH-TNF-α ARE complexes with respect to various concentrations of wild type LmGAPDH. Panel A (Lower panel), depicting % of the shifted band intensities of upper panel 1A with respect to various concentration of LmGAPDH. Panel B (Upper panel), represented the REMSA image for inhibition of TNF-α ARE binding of LmGAPDH by NAD^+^. Panel B (Lower panel), depicting % of the inhibition of shifted band intensities with increasing concentration of NAD^+^. Band intensity was quantified by ImageJ software (NIH). Background signals from blank regions of the nylon membranes were subtracted from the signal intensities obtained from shifted RNA. All data are representative of three independent experiments. Error bars represent the SD from three independent experiments.

The nature of the inhibition by TNF-α ARE was further investigated by activity measurement with increasing concentrations of NAD^+^ in presence of different concentrations of TNF-α ARE (Figure 2A). The double-reciprocal plots (Figure 2B) for GAPDH activity versus NAD^+^ in presence of varying concentrations of TNF-α ARE showed that the apparent Km value of NAD^+^ increases with increasing concentrations of TNF-α ARE without affecting the V_max_ value. These data indicate that the inhibition by TNF-α ARE is competitive in nature with respect to NAD^+^ binding.

**Figure 2.**
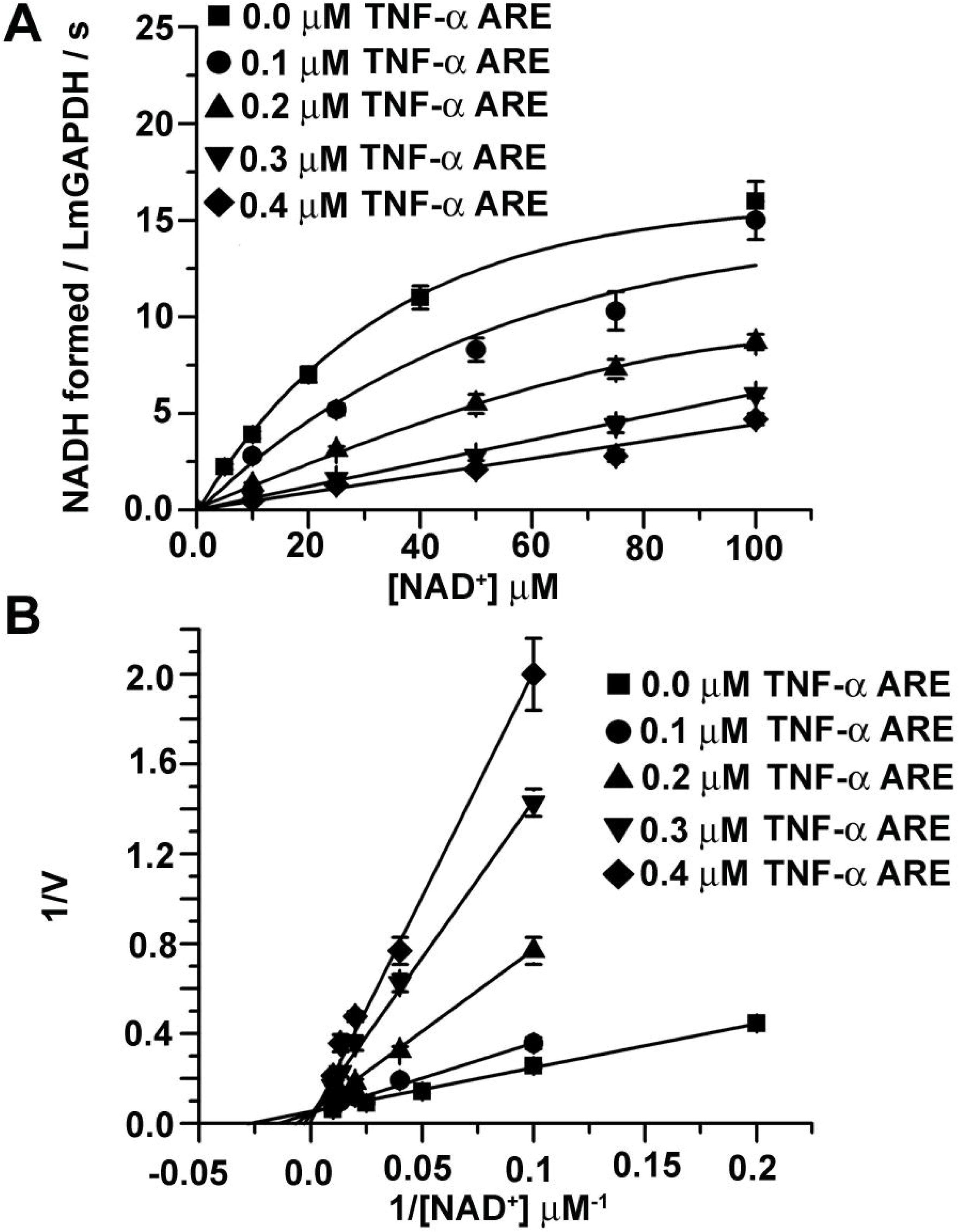
NAD^+^ dependent activity measurement in presence of different concentration of TNF-α ARE. Panel A denoted the turn over number of wild type LmGAPDH versus NAD^+^ concentration in presence of various concentration of TNF-α ARE (0-0.4 µM). Panel B represents the double-reciprocal plots for GAPDH activity versus NAD^+^ in presence of different concentrations of TNF-α ARE (0-0.4 µM). All data are representative of three independent experiments. Error bars represent the SD from three independent experiments.

### Homology modelling

As of now, NAD^+^ binding residues of GAPDH that are also responsible for TNF-α ARE binding is still unexplored. Similar to primary structure of human GAPDH, LmGAPDH is composed of two domains: the NAD^+^-binding domain (blue colored; residues 1–163 and 335-355) and the catalytic domain (orange colored; residues 164–334; Figure 3A). Homology alignment studies among human, bacterial and parasitic GAPDH suggest that the NAD^+^ binding site in LmGAPDH is conserved at position of 9-16 (NGFGRIGR), D39, 112-113 (TG), and N336 (Figure 3A). On the basis of the homology model, the structural features of LmGAPDH are found to be similar to those of *Leishmania mexicana* (Lmx) GAPDH [35] ( PDB entry code: 1a7k). Both NAD^+^ binding and catalytic domains contain eight β-sheets each (Figure 3B). These β-sheets are interconnected by either short loops or helices. Model structure of LmGAPDH covers 99.9% amino acid residues and sequence identity coverage is 94.69% with respect to reference X ray crystal structure of LmxGAPDH. Model was assessed with SWISS-MODEL server. Root mean square deviation (RMSD) value of our model structure from reference structure is very low (0.08 Å). Taking advantage of the coordinates of the model, we have searched for the NAD^+^ binding residues. Based on NAD^+^ bound LmxGAPDH crystal structure, our model structure shows that backbone amino groups of Arg13 and Ile14 at glycine-rich loop (residues 10–16) make hydrogen bonds with pyrophosphate moiety of the NAD^+^ (Figure 3C). The adenosine ribose moiety of NAD^+^ interacts with Asp39 residue of LmGAPDH via two hydrogen bonds. The Asn336 residue of LmGAPDH is hydrogen bonded to nicotinamide carbonyl oxygen. In addition, the model structure predicts that the distances from Arg16 and Thr112 residue to the NAD^+^ are 5.5 and 3.94 Å, respectively. In contrast to LmxGAPDH structure, bovine GAPDH and Group B *Streptococcus* GAPDH structure have shown that two water molecules bridge the glycine-rich loop and the cofactor pyrophosphate in the active site of the NAD^+^ bound complex [31, 32].

**Figure 3.**
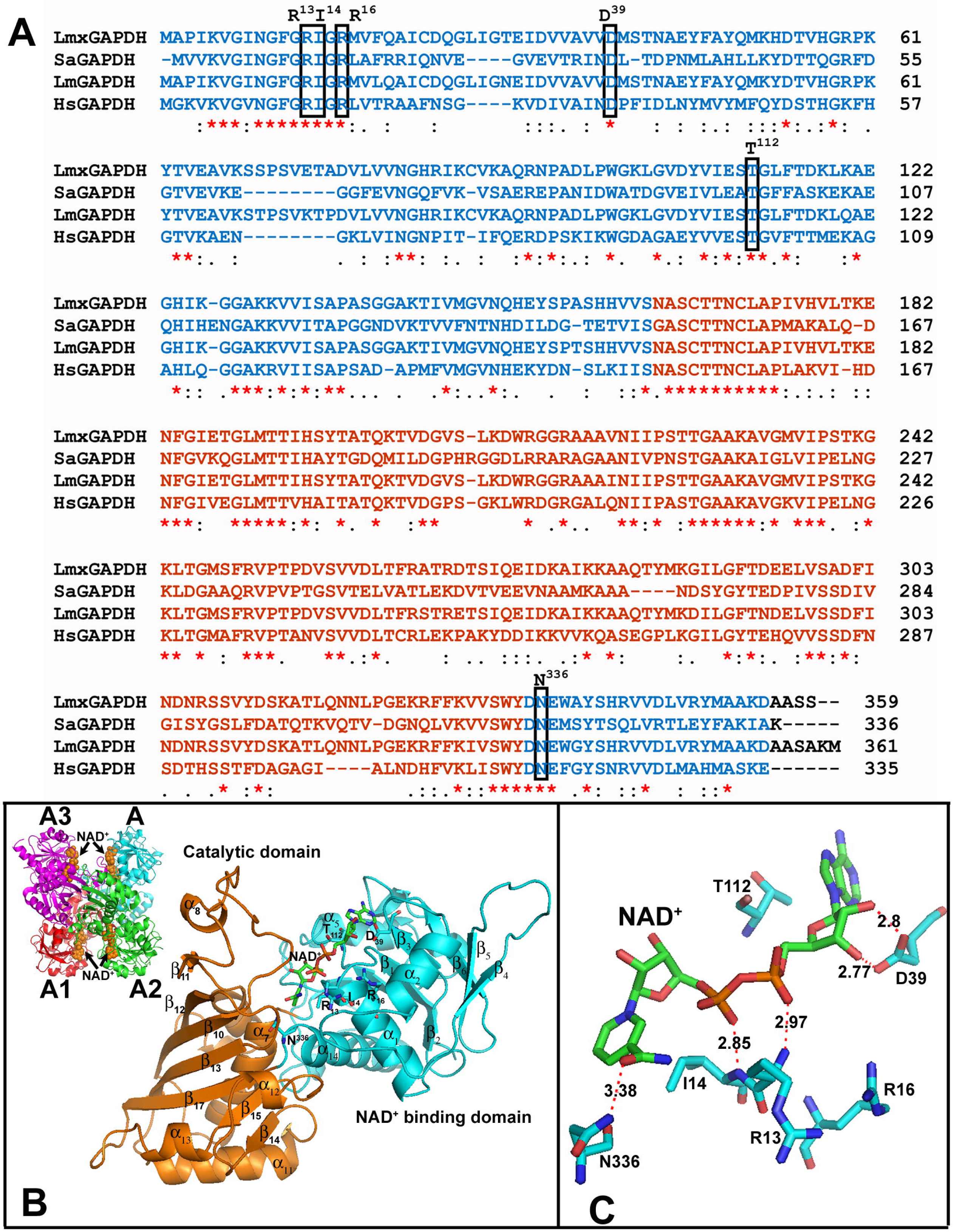
Sequence alignment and model structure of LmGAPDH. The sequence of LmGAPDH was aligned with the GAPDH from *Leishmania mexicana* (LmxGAPDH), *Streptococcus agalactiae* (SaGAPDH) and *Homo sapiens* (HsGAPDH). Blue and orange color denoted NAD^+^ binding domain and catalytic domain, respectively. * denoted identical residue. Panel B, based on the published X-ray crystallographic structures of *Leishmania mexicana* GAPDH (PDB entry code: 1a7k), we constructed a three-dimensional model of the LmGAPDH-NAD^+^ complexes by knowledge-based homology modeling using the SWISS-MODEL [51] and PyMOL software (http://www.pymol.org/pymol). The ribbon structure of NAD^+^-binding domain and catalytic domain are cyan and orange colored, respectively. Inset of panel B showed homo tetrameric three-dimensional model of the LmGAPDH-NAD^+^ complexes. The four homo subunits denoted A (cyan), A1 (red), A2 (green), and A3 (magenta). Panel C, the interaction between residue and NAD^+^ without ribbon.

### Expression and purification of the wild-type and mutant LmGAPDH enzymes

Since it has been experimentally observed that incubation of LmGAPDH with its cofactor NAD^+^ reduces its interaction with TNF-α ARE (Figure 1 and 2), we were interested in knowing whether NAD^+^ binding residues are involved at all in the TNF-α ARE binding. Based on the sequence analysis and structural studies, the following residues are present at or near NAD^+^ binding domain: R13, I14, R16, D39, T112 and N336 (Figure 3). For validating our hypothesis, the following LmGAPDH mutants were constructed: R13A, R13K, I14A, R16A, D39A, T112A and N336A. The wild-type and mutant LmGAPDH were overexpressed in *E. coli* and purified by using a Ni^2+^-NTA column as N-terminal six His-tagged proteins. All LmGAPDH mutants were soluble and purified for subsequent characterizations. The level of purity of all wild-type and mutant proteins were measured by running on a 12% SDS-polyacrylamide gel, followed by staining with Coomassie Brilliant Blue. All purified wild-type and mutant proteins were homogeneous and of similar molecular mass (Figs. 4A-B).

**Figure 4.**
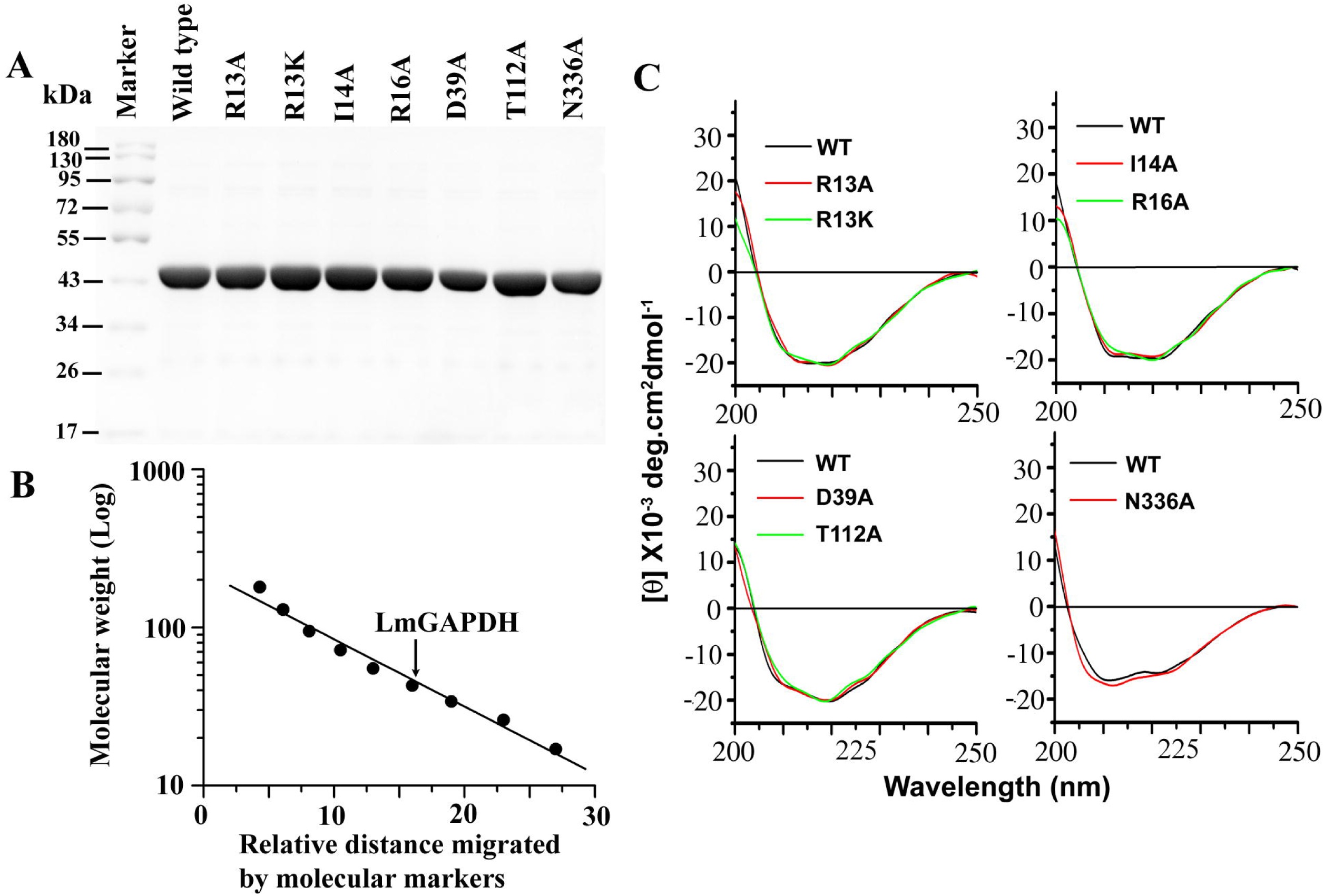
SDS-PAGE and CD analysis of different mutants. Panel A shows the level of purity of wild type and mutant proteins, which was determined by separation on a 12% SDS-polyacrylamide gel, followed by staining with Coomassie Brilliant Blue. Panel B shows molecular weight versus relative distance migrated by molecular markers. Panel C, shows CD spectra of wild type and mutant proteins. The results were expressed as molar ellipticity [ө] in units of m degree g cm^2^ dmol^-1^ versus wavelength.

### Circular dichroism (CD) spectroscopy measurement of wild type and mutant LmGAPDH

The characteristic helical secondary structure of folded proteins display the minima at 221 and 208 nm in the ultraviolet wavelength region (UV) of CD spectra [36]. To test whether LmGAPDH maintains its native helical-rich structure after mutation at several NAD^+^ binding residues, the UV-CD spectrometry was performed. UV-CD analysis (Figure 4C) suggested that the secondary structure of LmGAPDH protein is not significantly changed after point mutation at the position of 13, 14, 16, 39, 112 and 316. These results can rule out the possibility of inactivation being due to changes in the secondary structure of the enzyme after mutation.

### Kinetic properties of wild type and mutants LmGAPDH

To determine the kinetic parameters (Km for NAD^+^ and *k_cat_*) of wild type and mutants LmGAPDH, the enzymatic rate was spectrophotometrically measured at 340 nm in presence of different NAD^+^ concentration under steady-state conditions (Figure 5). Like *Trypanosoma cruzi* GAPDH [37], the wild type LmGAPDH catalyzed reactions exhibited NAD^+^ concentration dependence and followed saturation kinetics. Table 1 shows the comparison of the Km and *k_cat_* values of all seven mutants with wild-type protein at neutral pH 7.5. The GAPDH activity of the R13A, R13K and N336A enzymes were reduced to zero, hence their Km for NAD^+^ could not be determined. Interestingly, mutation of Arg-13, which forms hydrogen bonds with pyrophosphate moiety of the NAD^+^, to either lysine or alanine also showed complete loss of activity, suggesting that this Arg residue at position-13 is absolutely indispensable for enzymatic function of LmGAPDH. Since no improvement in the enzymatic turnover was observed for the R13K mutant, it is likely that the positive charge of the sidechain is not important for catalytic activity or NAD^+^ binding. It might be due to backbone amino group rather than positively charged guanidinium group of Arg13, which makes hydrogen bonds with pyrophosphate moiety of the NAD^+^. Another mutation of the invariant residue N336, which fixes the plane of the nicotinamidium ring of NAD^+^ by hydrogen bonding, to alanine showed completely inactive protein, indicating that the interaction between the nitrogen of Asn with the oxygen atom of the carboxyamide of the pyridinium ring is essential for catalytic function of LmGAPDH. To assess the significance of hydrogen bond between the backbone amino group of Ile-14 and pyrophosphate moiety of the NAD^+^ in cofactor binding, as proposed from the model structure, we have substituted this Ile-14 residue to an alanine by site-directed mutagenesis. The I14A mutant exhibited a significant amount of activity loss with ∼ 97% reduction in *k_cat_* value with compared to the wild-type enzyme. This mutant also showed the alteration of Km value for NAD^+^. Together, all data suggest that the involvement of Ile-14 residue is a crucial for catalysis as well as binding of the cofactor NAD^+^. To evaluate the importance of two hydrogen bonds between the side chain carboxy group of Asp-39 and 2, 3-hydroxy groups of adenosyl ribose of NAD^+^ in cofactor recognition, as proposed from the model structure, we have replaced this Asp-39 residue to a neutral alanine by site-directed mutagenesis. The Km value of D39A for NAD^+^ binding is 12-fold higher than the wild type protein suggesting that these H-bond interactions between D39 and NAD^+^ play an important role for NAD^+^ binding. The catalytic activity of the D39A mutant showed only ∼11% enzymatic activity with respect to wild type protein indicating that this acidic residue is crucial for catalytic function of LmGAPDH. In addition, we measured the kinetic parameters of other two mutants (R16A and T112A) those residues which are present near the NAD^+^ binding site. The apparent Km values of both the R16A and T112A mutants for NAD^+^ showed an approx. 4-fold higher values as compared with the wild type, subsequently *k_cat_* values of these mutants reduce to 50%. All these results reaffirmed the importance of these highly conserved residues in the NAD^+^ binding site that had been suggested by crystallographer [30].

**Figure 5.**
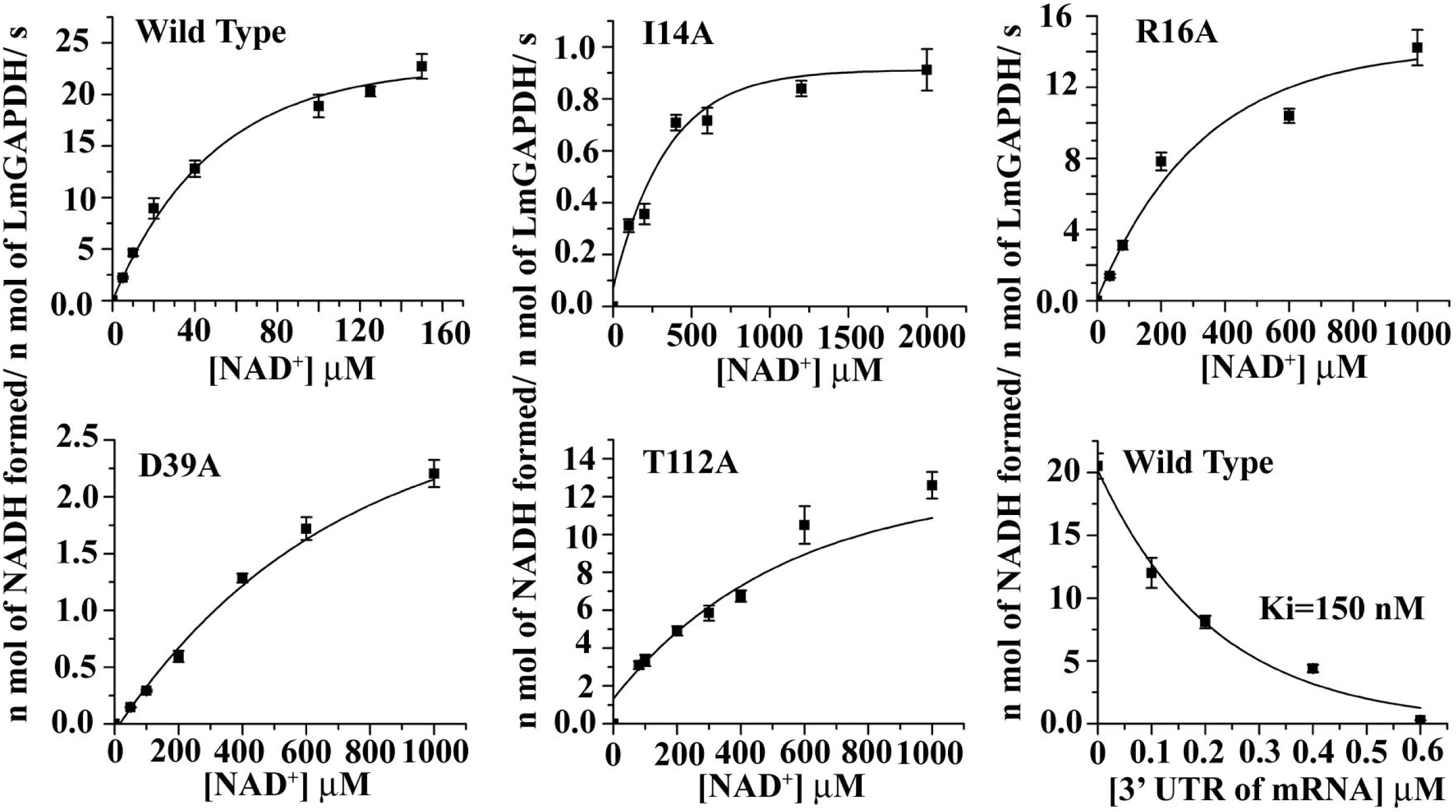
NAD^+^ dependent enzymatic activity of wild type and mutants LmGAPDH. Velocity of wild-type, I14A, R16A, D39A, and T112A proteins were plotted against varying concentrations of NAD^+^. Last panel shows the velocity of wild type protein in presence of saturated concentration of NAD^+^ versus TNF-α ARE concentration. All data are fitted to a first order curve. The Km and *k_cat_* values from the fitting curve are listed in Table 1. Data were plotted as means and SD from three independent experiments for each assessment.

**Table 1.** Comparative study for GAPDH activity and Km value among different mutant proteins. The catalytic activities were determined at 25 °C as described under “Materials and Methods.” The values represent the mean and standard deviation for three measurements each.

| Enzyme | Catalytic activity parameters |  |
| --- | --- | --- |
| | $K_m$ (μM) | $k_{cat}$ ($s^{-1}$) |
| Wild type | $48 \pm 3$ | $24.8 \pm 1.1$ |
| R13A | Not measuarable | No activity |
| R13K | Not measuarable | No activity |
| I14A | $340 \pm 15$ | $0.8 \pm 0.1$ |
| R16A | $380 \pm 43$ | $14.2 \pm 1.8$ |
| D39A | $595 \pm 78$ | $2.8 \pm 0.1$ |
| T112A | $423 \pm 44$ | $11.2 \pm 0.6$ |
| N336A | Not measuarable | No activity |

### Comparison of TNF-α ARE binding property of the wild-type with that of the mutant enzymes

Experimental observations from above results have shown that TNF-α ARE acts as an inhibitor of LmGAPDH by competing for the NAD^+^-binding site (Figs. 1 and 2). Several studies have suggested that RNA binding is mediated by the NAD^+^ binding domain [25, 28, 38]. We were interested to find out whether NAD^+^ binding residues are involved in the TNF-α ARE-binding. If NAD^+^ binding residues are part of the RNA binding site, then the binding of the TNF-α ARE to the enzyme could be hampered by mutation at NAD^+^ binding site. These effects can be quantified by the measurement of shift band in RNA electrophoretic mobility shift assay. Thus, it would be rational to draw conclusions that any alteration in the TNF-α ARE binding affinity of the mutant enzymes should be echoed in its dissociation constant (K_D_) value. To investigate whether the mutations cause any change in the TNF-α ARE-binding performance (K_D_ value), we measured % shifted band intensity with respect to different concentration of mutant proteins. Figure 6 shows a comparative study of the amount of gel shifted band among different mutant enzymes. Panel A shows that the same amount of the various purified mutant enzymes were incubated separately with TNF-α ARE in REMSA assay mixture under conditions where maximum amount of the wild-type LmGAPDH (50 µM) binds. In case of R13A, R13K and N336A enzymes, we did not get any detectable amount of the gel shifted band. It is clear from these results that the R13A, R13K or N336A mutants did not bind to the TNF-α ARE at all, implicating absolute indispensable of these residues in TNF-α ARE-binding. The results from R13A and R13K mutants suggest that not only the positive charge in the side chain but also the specific orientation with size of this Arg13 residue are strictly necessary for suitable binding of the TNF-α ARE. Like R13 residue, the bonding network between N336 and TNF-α ARE is strictly restricted for the TNF-α ARE-LmGAPDH complex formation. On the other hand, there were significant alteration in the TNF-α ARE-binding property of the I14A, R16A, D39A or T112A mutant proteins. To measure the K_D_ value of mutant LmGAPDH protein for TNF-α ARE, we plotted the % of shifted band intensity of mutant protein-RNA complexes versus the concentration of mutant LmGAPDH (Figure S1). Enzyme dependent TNF-α ARE-binding did not occur in case of R13A, R13K and N336A mutants (Panel B, C and H), hence their K_D_ value for TNF-α ARE binding could not be determined. Although a very low amount of shifted band was observed in I14A but the band intensity of I14A-TNF-α ARE did not change significantly with increasing concentration of mutant protein (Panel D) indicating that TNF-α ARE binding to I14A mutant was too weak to be reliably determined by REMSA. These data suggest that the participation of Ile-14 residue is crucial in the LmGAPDH-TNF-α ARE complex formation. The K_D_ value of the D39A LmGAPDH for TNF-α ARE (21 ± 1.8 µM) was increased ∼12.5 fold compared to the wild type protein (∼1.7 µM) suggesting that the side chain carboxy group of Asp-39 residue may play an important role for the interaction with adenosyl ribose moiety of TNF-α ARE. In addition, we also measured the K_D_ value of other two mutants (R16A and T112A) for TNF-α ARE, which are located near the NAD^+^ binding site. The K_D_ value of R16A and T112A LmGAPDH protein for TNF-α ARE were 4.6 ± 1.1 µM, and 8.6 ± 1.2 µM respectively. Thus, the K_D_ value of the R16A and T112A LmGAPDH for TNF-α ARE were increased by ∼2.6 and 5-fold compared to the wild type protein, respectively. Altogether, above findings suggest that NAD^+^ binding residues are involved in the TNF-α ARE-binding.

**Figure 6.**
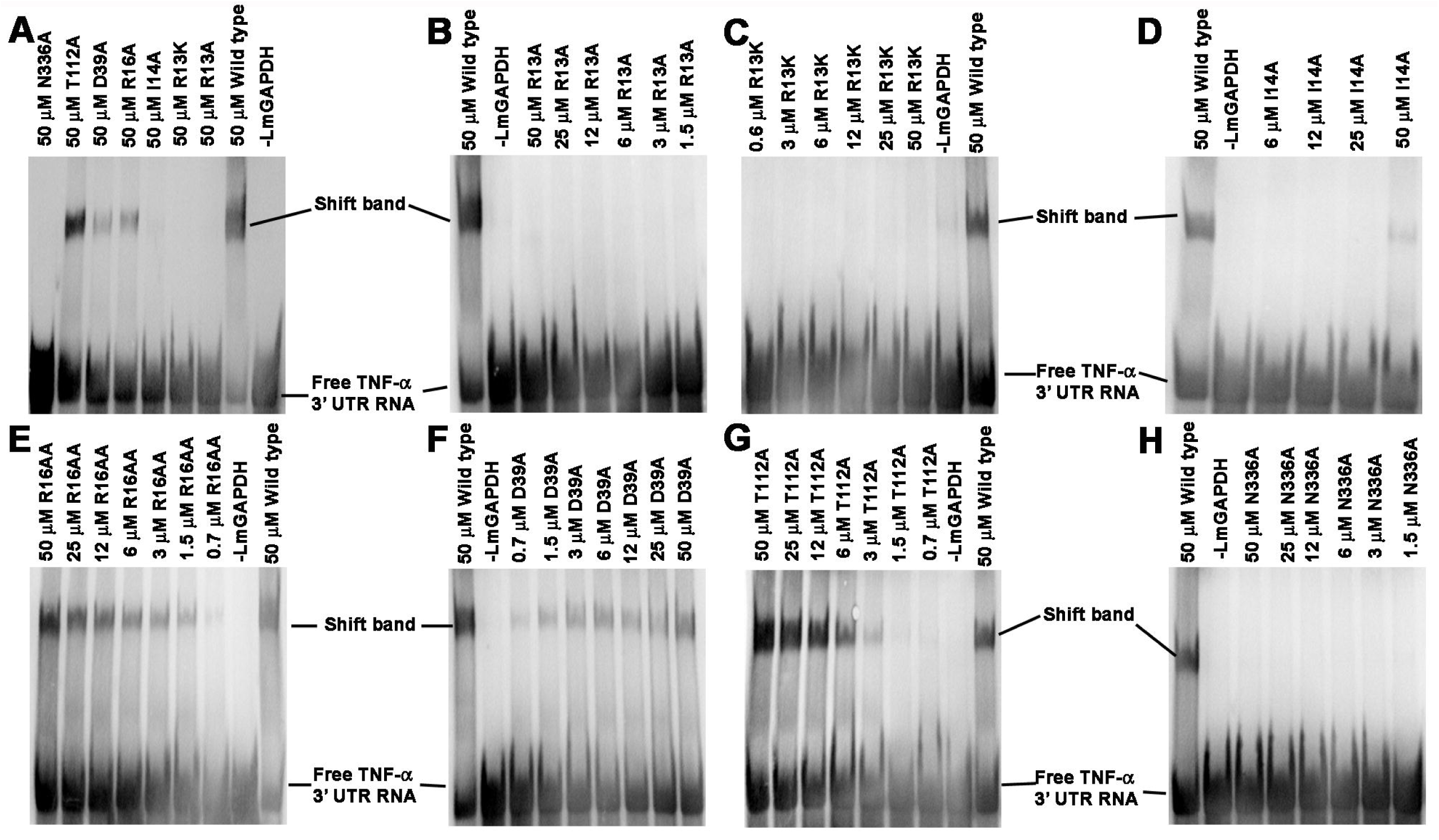
Comparative study among different mutant proteins for TNF-α ARE binding by REMSA. Panel A, the REMSA measurement for wild type (WT) and mutants LmGAPDH binding to the TNF-α ARE. Panel B, C, D, E, F, G and H denoted the REMSA image for the binding pattern of TNF-α ARE towards different mutant proteins with respect to various concentrations of R13A, R13K, I14A, R16A, D39A, T112A and N336A mutants, respectively. All data are representative of three independent experiments.

To confirm the K_D_ values for TNF-α ARE obtained by REMSA, protein induced fluorescence enhancement (PIFE) was performed for measuring the interactions of enzyme with TNF-α ARE at pH 7.5 (Figure 7). We measured the steady-state binding affinity (K_D_ values) of immobilized Cy3 labeled TNF-α ARE probe (Figure 7A, Scheme of TNF-α ARE probe) with an unlabeled LmGAPDH protein by PIFE in a microwell plate (mwPIFE). Interestingly, the mwPIFE values with LmGAPDH showed a highly significant difference from control reactions with bovine serum albumin (BSA) demonstrating that mwPIFE can reliably measure a protein-RNA complex formation in a quantitative manner. In contrast to wild type protein, the mwPIFE value of R13A, R13K, I14A or N336A mutants did not show any fluorescence enhancement, implicating absolute necessity of these residues in TNF-α ARE-binding (Figure 7B). The mwPIFE value of the fluorescent TNF-α ARE probe was dramatically enhanced by increasing concentration of wild type LmGAPDH, whereas mwPIFE value of the fluorescent TNF-α ARE probe with the R16A, D39A and T112A mutant LmGAPDH increased only slightly at the highest concentrations used in the assay (Figure 7C). The LmGAPDH dependent mwPIFE data were best fitted to saturation kinetics. PIFE measurement suggest that the calculated K_D_ value of wild type LmGAPDH for TNF-α ARE is 1.8 µM, which is similar with the K_D_ values obtained by REMSA (1.7 µM). The K_D_ value of R16A, D39A and T112A LmGAPDH protein for TNF-α ARE were 7.77 ± 0.62 µM, 9.53 ± 0.8 µM and 6.68 ± 0.55 µM, respectively, those are within the range previously determined by REMSA experiments. Altogether, these data further prove that NAD^+^ binding site residues play an important role for TNF-α ARE binding.

**Figure 7.**
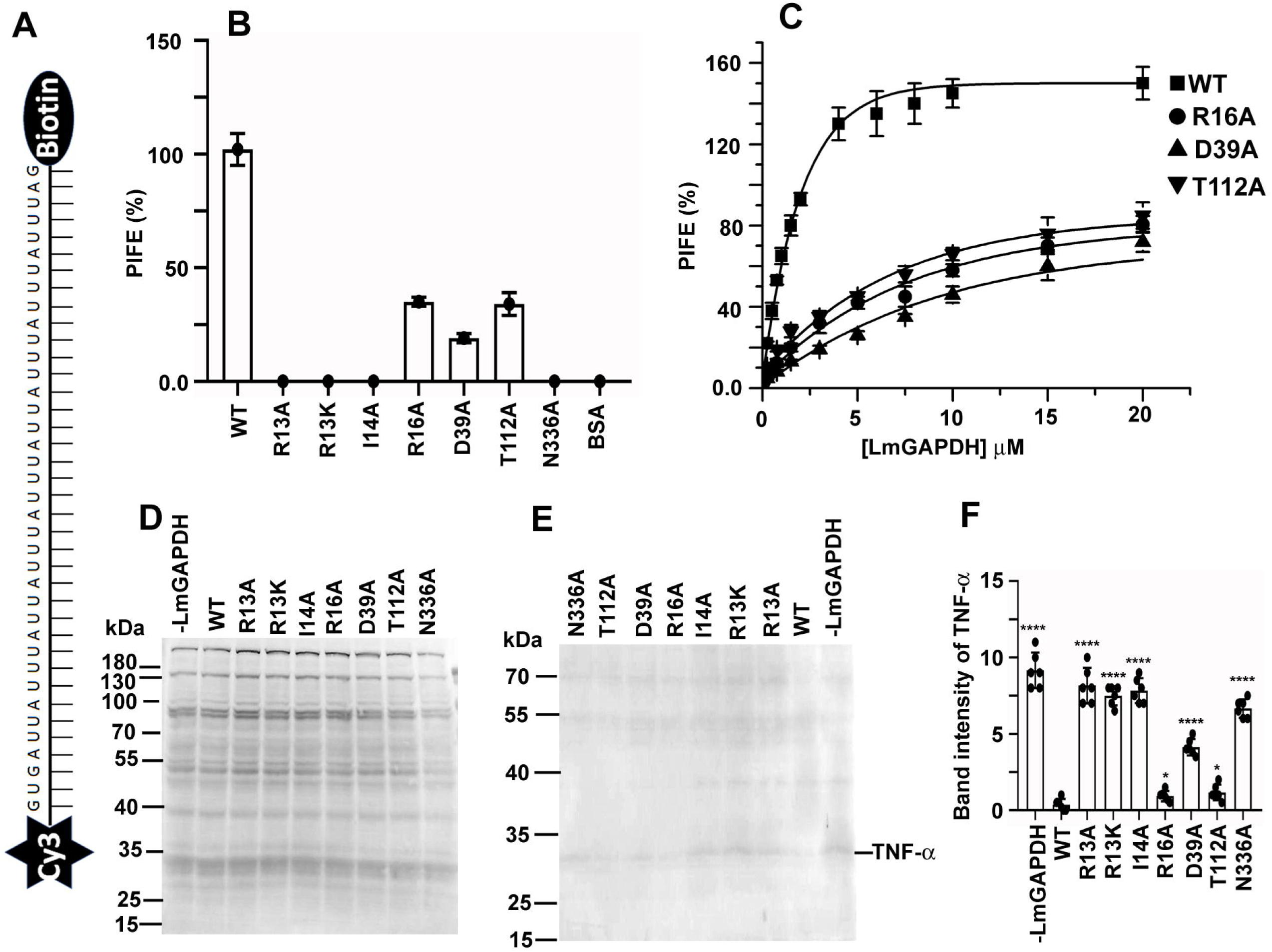
Analysis of LmGAPDH and TNF-α ARE interaction by mwPIFE and *in vitro* protein translation. A, Scheme of TNF-α ARE probe for detecting LmGAPDH binding. B, Mutation on LmGAPDH affects the magnitude of mwPIFE. The concentration of wild type as well as mutant protein used was 4.0 μM. Undetectable PIFE value observed in case of R13A, R13K, I14A, N336A and BSA protein. C, LmGAPDH concentration dependent mwPIFE value. Error bars indicate standard deviations from measurements of three independent wells. Calculated K_D_ values for each enzyme are: wild type (1.8 ± 0.11 μM), R16A (7.8 ± 0.6 μM), D39A (9.5 ± 0.8 μM), T112A (6.7 ± 0.6 μM). D, the image of *in vitro* general translation pattern in presence of wild type and mutant protein. E, the image of *in vitro* TNF-α translation pattern in presence of wild type and mutants of LmGAPDH. Panel F, depicting the band intensities of TNF-α expression after normalization with the background in presence of various GAPDH mutants. Band intensity was quantified by ImageJ software (NIH). All data are representative of six independent experiments. Error bars represent the SD from six independent experiments. Statistical analysis for parametric data was calculated by Student’s t test or analysis of variance (ANOVA). **** and * denoted p value of less than 0.0001 and 0.05, respectively.

To reconfirm the REMSA and mwPIFE data, *in vitro* protein translation was performed for measuring the interaction of enzyme with TNF-α mRNA. Earlier, *in vitro* protein translation results suggested that the LmGAPDH-TNF-α mRNA complexes inhibit translation of TNF-α expression [20]. If the mutations present in NAD+ binding site bring down the binding ability of TNF-α ARE with LmGAPDH then these mutations would show higher amount of TNF-α protein synthesis compared to wild type protein. To test this hypothesis, we investigated *in vitro* protein translation of TNF-α in the presence of equal amount of wild type or mutant LmGAPDH (Figure 7D-E). The *in vitro* general translation band pattern in all lanes is similar (Figure 7D), suggesting that protein synthesis occurred in the presence of wild type or mutant LmGAPDH. After immunoprecipitation with TNF-α monoclonal antibody, the blot showed that the TNF-α expression is inhibited in presence of wild type and T112A mutant LmGAPDH (Figure 7E). In contrast to wild type protein, R13A, R13K and N336 mutant proteins could not inhibit *in vitro* TNF-α protein synthesis suggesting that these mutant proteins are unable to bind with TNF-α mRNA. Like binding capacity of mutants, the order of *in vitro* protein synthesis band was R16A <D39A<I14A. Altogether, these data further reconfirm that NAD^+^ binding site residues play a vital role for TNF-α ARE binding.

## Discussion

Combined use of systematic mutational analysis and homology modelling guided us to pinpoint the key active site residues for cofactor NAD^+^ binding and also establish the novel TNF-α ARE interaction with NAD^+^ binding residues of LmGAPDH. Despite having structural differences between NAD^+^ and mRNA of TNF-α, both occupy a closely overlapping position of the enzyme with similar interacting residues has provided important mechanistic insights into its catalytic process and post transcriptional regulation of TNF-α mRNA [20]. In this article, we have shown for the first time that cofactor NAD^+^ binding residues of LmGAPDH are crucial for TNF-α ARE binding. Here, we discuss the following salient points: (a) Binding studies and enzymatic activity measurement have shown that the TNF-α ARE is competitively inhibited by the cofactor NAD^+^ binding in LmGAPDH; (b) We have demonstrated that R13 and N336 mutation at the NAD^+^ binding site of LmGAPDH generate a catalytically dysfunctional mutant-isoforms those are unable to bind TNF-α ARE; (c) Furthermore, other I14A, R16A, D39A and T112A mutations in NAD^+^ binding site showed higher Km values (lower binding affinity) for cofactor NAD^+^ as well as higher K_D_ values for TNF-α ARE compared to the wild-type protein; (d) Both I14A and D39A single mutants were affected more drastically on the catalytic activity as well as the TNF-α ARE binding compared to R16A and T112A mutations.

In silico model of LmGAPDH demonstrates that the backbone amino groups of Arg13 and Ile14 at glycine-rich loop (residues 10–16) form hydrogen bonds with pyrophosphate moiety of the NAD^+^. These two residues are located far apart from the substrate-glyceraldehyde-3-phosphate binding region, which may avoid any unwanted impact on substrate-binding. Our findings demonstrate that a point mutation at the Arg13 is sufficient to knock out the catalytic capacity of GAPDH due to loss of cofactor NAD^+^ binding. Substitution of Ile14 residue by Ala caused ∼97% activity loss and a significant increase in the Km value for NAD^+^, thereby influencing substantial weakening of the cofactor binding. Our findings thus amply corroborate an earlier study with human GAPDH, which also concluded that a mutation at analogous arginine (R13) and isoleucine (I14) in the NAD^+^-binding site is sufficient to generate a dysfunctional GAPDH [39]. Our prediction was that NAD^+^ binding residues might be involved in TNF-α ARE binding due to competitive inhibition. Upon alanine replacement, the R13A mutant was influenced more drastically (not any TNF-α ARE binding) than the I14A mutant that displayed considerably reduced, but detectable TNF-α ARE binding. The higher impact of the Arg13 compared to the Ile14 residue reveals the variations in the strengths of H-bond between the respective backbone amino groups and pyrophosphate moiety of the NAD^+^, possibly due to differing H-bond length and influence of water molecule. These results definitely suggest that both Arg13 and Ile 14 are indeed crucial TNF-α ARE binding residues. Furthermore, this specific arginine residue of LmGAPDH is so restricted to replacement that its substitution, even with a lysine residue also recognized to retain the similar charge in the side chain, led to loss of cofactor NAD^+^ or TNF-α ARE binding. These results endorsed us to suggest that the positive charge of guanidinium group in the side chain, the shape and size of this Arg13 residue are compulsory for proper interaction of the cofactor NAD^+^ or TNF-α ARE, thus providing a justification for the firm preservation of the amino acid among other GAPDH including human, bacteria and parasites.

The position of Asp39 residue relative to the bound NAD^+^ predicted the bidentate hydrogen-bond formation between the 2′- and 3′-hydroxy groups of adenosine ribose moiety of NAD^+^ and the carboxy group of Asp39. Replacement of this residue by alanine caused a substantial enhancement in the Km value for NAD^+^, thereby influencing significant destabilization of the cofactor binding to the LmGAPDH. Therefore, based on combination factors of both the structural analysis and the kinetic data, the present study suggests that the formation of bidentate hydrogen-bond between the -COOH group in the sidechain of Asp39, and the 2′-& 3′-OH groups of the adenosine ribose is one of the key influential features for the LmGAPDH-NAD^+^ complex formation. Therefore, it is possible that TNF-α ARE may bind in or close to the Asp39 residue in NAD^+^ binding site. The mutant displayed ∼12.5-fold lower affinity towards TNF-α ARE binding suggesting that the ribose binding site is also important for proper interaction with the TNF-α ARE.

Another evidence for this model-based prediction was achieved while experimental observation was shown that (i) the N336A protein is catalytically inactive, and (ii) substitution of N336 by neutral alanine caused in non-binding nature of the enzyme with TNF-α ARE. The TNF-α ARE - binding property of Asn336 documented in the present study is hitherto unknown and thus provided an attractive point of the enzyme-NAD^+^ interaction as well as enzyme-TNF-α ARE binding. Earlier, replacement of Asn313 (analogous of Asn 336 of LmGAPDH) in the *Escherichia coli* GAPDH by Thr or Ala residue indeed caused ∼50 fold less activity and a significant increase in the Km value for NAD^+^, thereby influencing substantial weakening of the cofactor binding [40]. It is clear from X-ray crystal structure that N313T mutant GAPDH feebly interacts with NAD^+^ compared to the wild-type enzyme [41]. This particular Asn residue had previously been proposed from their crystal structure that one of the structural factors contribute to fix in a productive manner the plane of the nicotinamidium ring of NAD in a syn orientation by hydrogen bonding between its nitrogen ND2 and the oxygen of the carboxamide at the C-3 atom of the pyridinium [42]. As the lacking of nicotinamide nucleotide in TNF-α ARE, the mechanism of its binding by Asn336 is still not known. One plausible explanation could account for this paradoxical result that is Asn336 might form H-bond with other nucleotide base (uracil) in TNF-α ARE instead of nicotinamide. Further x-ray crystal structure of GAPDH and TNF-α ARE complexes are needed to confirm interaction site of TNF-α ARE with Asn336.

Kinetic studies from our laboratory showed that TNF-α ARE was a competitive inhibitor of GAPDH with respect to NAD^+^, having Ki values in the range of 150 nM level (Figure 5). It has been suggested that inhibitory effect by TNF-α ARE is via competitively occupying the NAD^+^-binding site. Our mutational study at the adenosine pyrophosphate moiety and adenosine ribose site, and nicotinamide binding residues in the NAD^+^-binding site further suggest that NAD^+^ binding residues are responsible for TNF-α ARE binding. Earlier studies have suggested that the AU-rich RNA binding site may be localized to the two positively charged NAD^+^ binding groove of the GAPDH tetramer [43]. They hypothesized that one tetrameric GAPDH might be interacting with two separate RNA strands or two separate AREs in a single RNA strand. From our structural analysis, we found that the NAD^+^ molecule is localized on the positively charged surface of LmGAPDH (Figure 8A). The residues Gly12, Arg13, Ile14, Gly15, Arg16, Asp39, Arg93, Ser135 and Asn336 tightly encircled the dineucleotide ring of NAD^+^ (Figure 8A). Because of a common adenine base, phosphate moiety and ribose base, AU-rich RNA may be bound tightly to this positively charged surface groove of LmGAPDH (Figs. 8B and 8C). Interestingly, earlier researchers have shown from some instances that the RNA-binding activity of GAPDH depend on the glycolytic state of the cells [44]. Furthermore, they have shown that the metabolism shifts from oxidative phosphorylation to glycolysis in T cells during activation. At that time GAPDH prefers to engage with NAD^+^ binding rather than IFN-γ AREs binding, thereby allowing IFN-γ translation. Our previous *in vitro* protein translation data suggest that the TNF-α protein expression decreases with increasing LmGAPDH concentration [20]. Current *in vitro* protein translation results further reconfirm that NAD^+^ binding site residues play a vital role for TNF-α synthesis. In addition, we have demonstrated that the extracellular vesicles-derived LmGAPDH from *Leishmania* promastigotes selectively inhibited host TNF-α production. During *Leishmania* infection, the metabolic reprogramming is found to occur in parasite infected human macrophage as a result of increased oxidative phosphorylation relative to glycolysis [45]. At that moment, LmGAPDH seems to engage with host TNF-α mRNA binding rather than NAD^+^ binding, thereby inhibiting host TNF-α translation. Comparison of the amino acid sequences of GAPDH from prokaryotic to eukaryotic organisms suggests that mRNA-binding could be evolutionarily conserved. Thus, evolutionarily conserved role of GAPDH in mRNA-binding, possibly depending on the metabolic reprogramming of cells remains to be determined.

**Figure 8.**
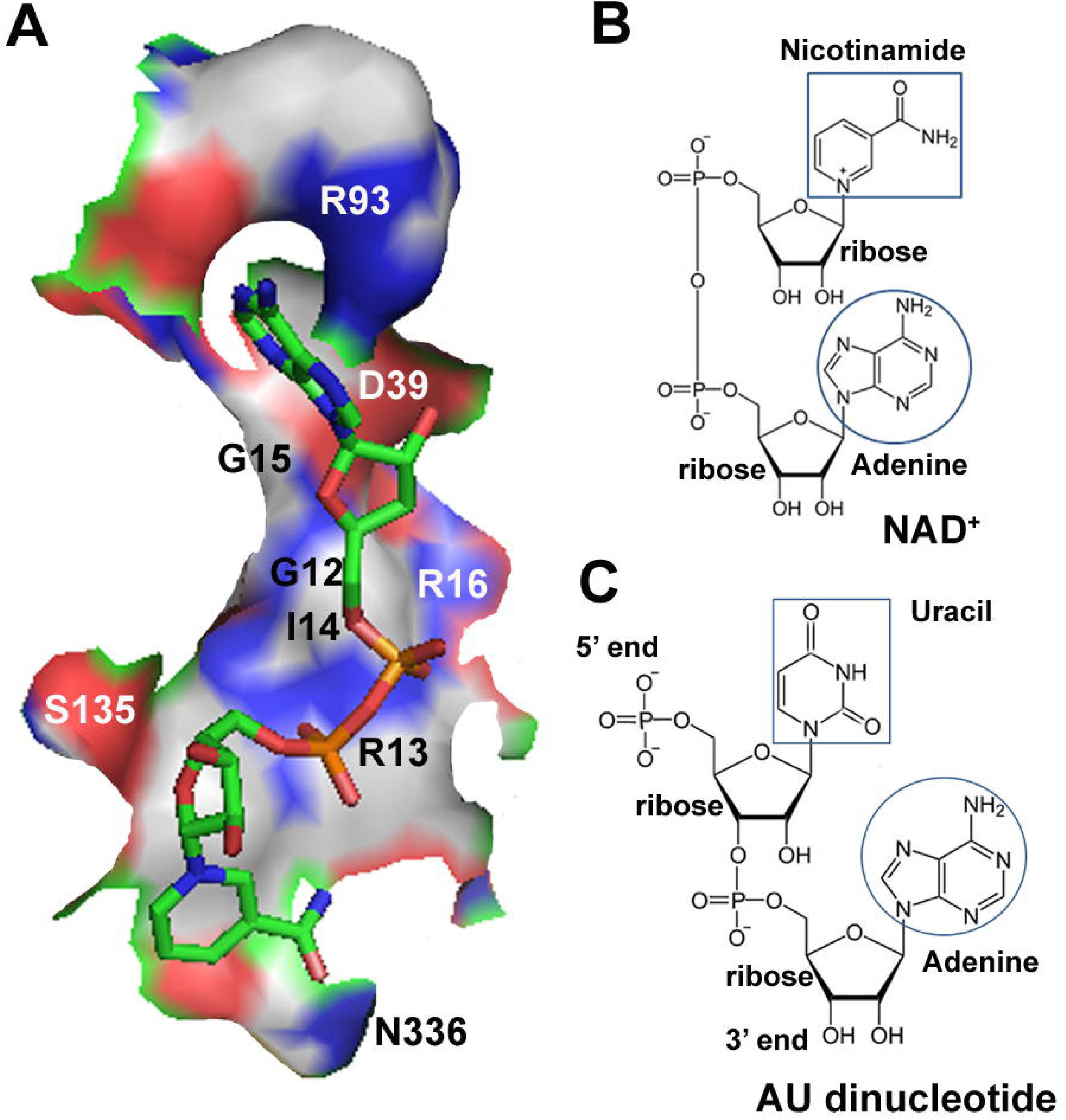
NAD^+^ interacts with LmGAPDH. Panel A, NAD^+^ molecule is localized on the positively charged surface of LmGAPDH. The residues Gly12, Arg13, Ile14, Gly15, Arg16, Asp39, Arg93, Ser135, and Asn336 tightly encircled the dineucleotide ring of NAD^+^. Panel B and C denoted the structural formula of nicotinamide adenine dinucleotide (NAD^+^) and AU (di-ribonucletide), respectively.

### Experimental procedures Materials

Imidazole and Ni^2+^-nitrilotriacetate resin were purchased from Sigma-Aldrich (St. Louis, MO, USA). All chemicals were procured from Sigma or sources reported earlier [46–48].

### Mutagenesis

Site-directed mutagenesis of wild type LmGAPDH in the pET15b was performed using the QuikChange site-directed mutagenesis kit from Agilent (Santa Clara, CA, USA). Mutations (underlined) and their corresponding oligonucleotides are mentioned in supporting information Table S1. The validation of mutations was checked by DNA sequencing.

### Expression and Purification of mutant LmGAPDH

The method for expression and purification of wild type LmGAPDH were described in our earlier paper [20]. In case of mutant proteins, pET15b vector containing mutant LmGAPDH constructs were transformed into *Escherichia coli* BL21(DE3) and were grown overnight in 50 ml of Luria-Bertani broth containing 200 µg/ml ampicillin at 37 °C temperature. This 50 ml overnight grown culture was then inoculated in 0.5 liter of terrific broth containing 200 µg/ml ampicillin at 37 °C temperature. When the culture reached an absorbance of ∼ 0.5-0.8 at 600 nm, 0.5 mM isopropyl β-D-1-thiogalactopyranoside was added and bacterial culture was further grown for ∼16 hours at 28 °C. Cells were then harvested by centrifugation at 6000 rpm for 10 min and washed 2 times with 1X phosphate-buffered saline. The pellet was resuspended in lysis buffer [50mM Tris-HCl (pH-7.5), 150 mM NaCl, glycerol (10%), β-mercaptoethanol, lysozyme (1 mg/ ml) and PMSF (4mM)]. The complete cell lysis was performed by sonication (Ultrasonic homogenizer, model-U250, Takashi) for 20 second pulse and 40 second gap repeated for 20 times in ice, followed by centrifugation at 14000 rpm for 1 hour. The supernatant was then loaded onto a pre-equilibrated Ni^2+^-NTA column which was then washed with equilibrium buffer [50 mM Tris-HCl (pH-7.5), 150 mM NaCl, glycerol (10%)] and with the same buffer containing 50 mM imidazole. The protein bound to the column was eluted with equilibrium buffer containing 250 mM imidazole. For removal of imidazole from the eluted protein, dialysis was performed. As expected, the purified wild type and seven mutant proteins showed a single protein band of approximately 39 kDa. The coomassie brilliant blue gel stained did not detect any other protein band except LmGAPDH band. The protein concentrations were measured by Bio-Rad Reagent with BSA as the standard.

### Enzyme assay using UV-visible spectroscopy

By using Shimadzu UV-2550 optical spectrophotometer, LmGAPDH activity was measured by monitoring NADH generation at 340 nm wavelength at 25°C. The assay mixture was composed of 40 mM Tris-HCl (pH-7.5), 50 mM K_2_HPO_4_, 2.5 mM D-glyceraldehyde-3-phosphate with an increasing concentration of NAD^+^. After incubating in a quartz cuvette of 1.0 cm path length for 5 minutes to attain temperature equilibrium and determine blank values, the reaction was initiated with the addition of respective 25 nM enzyme. Absorbance was recorded at 340 nm from 0 to 5 minutes. The molar extinction coefficient of NADH was 6.2 × 10^3^ M^-1^ cm^-1^ at 340 nm. For the determination of Km value for NAD^+^, we measured initial velocity of proteins at different concentrations of NAD^+^ at saturation level of glyceraldehyde 3-phosphate concentration (2.5 mM). The enzyme-catalyzed reactions in LmGAPDH exhibited saturation kinetics. The turnover number (*k_cat_*) and the Michaelis–Menten constant (Km) values were determined by using Michaelis–Menten enzyme kinetics in scientific statistics software (analysis of nonlinear regression) GRAPHPAD PRISM 6, GraphPad Software, Inc. CA, USA.

### RNA Electrophoretic mobility shift assay (REMSA)

RNA-EMSA was performed with the LightShift™ Chemiluminescent RNA EMSA kit (Thermo Scientific) according to the manufacturer’s instructions with slight modification. Briefly, an AU-rich RNA derived from TNF-α 3’ UTR (ARE 38-5’-GUGAUUAUUUAUUAUUUAUUUAUUAUUUAUUUAUUUAG-3’) was synthesized and biotin-labeled at 3’ end by Integrated DNA Technologies (IDT). The binding reaction involved 2 nM of biotin-labeled RNA probe along with respective purified LmGAPDH enzyme (wild type or mutant) in REMSA binding buffer (1X) with addition of 5% glycerol. The total reaction volume made up to 20µl; were incubated at room temperature for 30 minutes. Samples were separated on 6% native polyacrylamide gel in 0.5X Tris-Borate-EDTA (TBE) buffer at 100V, 4°C which were pre-run for 30 minutes. The gel was then transferred to positively charged nylon membrane (Sigma) using Trans-Blot SD semi-dry transfer cell (Bio-Rad) at 25V for 35 minutes. The biotin labeled RNA on the membrane was then cross-linked by a UV-light crosslinker (UVP HL-2000 HybriLinker™) and finally detected using streptavidin-horseradish peroxidase conjugate. Band shifts were visualized by exposure on iBright Imaging system (Invitrogen).

For determination of the K_D_ value of LmGAPDH protein for TNF-α ARE, we measured the % shifted band intensity with respect to different concentration of protein [49]. An exact value for the shifted RNA band intensity was determined by using ImageJ software. Background signals from blank regions of the nylon membranes were subtracted from the signal intensities obtained from shifted RNA. The fraction of RNA bound in each sample versus the concentration of LmGAPDH was plotted. Data from three independent experiments were analyzed using Prism software to perform non-linear regression with ‘One Site Specific Binding’ model {Y = B_max_*X/(K_D_ + X)} where Y is the band intensity, B_max_ is the fraction bound at which the data plateaus, X is the LmGAPDH concentration, and K_D_ is the apparent equilibrium dissociation constant.

### Protein induced fluorescence enhancement in a microwell plate (mwPIFE)

Protein induced fluorescence enhancement in a microwell plate (mwPIFE) was measured following the method described by Valuchova et al. [50] with slight modifications. Briefly, Pierce™ NeutrAvidin™ coated black 96-well plates (Thermo Scientific) were used for our mwPIFE experiments. The RNA probe, ARE-38 was synthesized and labeled at 5’ end with Cy3 fluorophore and at 3’ end with biotin by Integrated DNA Technologies (IDT). The wells were initially washed thrice with 200 µl of buffer containing 35 mM Tris-HCl (pH-7.5), 5.5% glycerol, 150 mM KCl, 1.0 mM EDTA, 0.1mM DTT and 1.5 mM Imidazole. 2.5 pmol of RNA probe was added to 100 μl of buffer and incubated for 2 hours in dark. The first scan was recorded after adding 100 μl of buffer in each well on Synergy H1 plate reader (BioTek), with excitation at 540 nm and emission at 580 nm. The previous buffer was replaced with buffer containing the respective LmGAPDH protein (wild type, mutant or BSA). Incubations were carried out at room temperature for 30 minutes to reach binding equilibrium and the second scan was recorded. %PIFE = [{(2^nd^ scan-1^st^ scan)/1^st^ scan}X100].

### Circular dichroism measurement

The circular dichroism spectra were obtained in a MOS-500 Circular dichroism spectrometer of BioLogic Science Instruments. 10.0 µM wild type or mutant LmGAPDH protein in buffer containing 10 mM Tris-HCl (pH-7.5) and 50 mM NaCl were purged with nitrogen and CD measurement was conducted from 200 to 250 nm. All the measurements were recorded at room temperature in a quartz cuvette using a cell with a 1-mm light path, averaging 10 scans per sample with 1 nm step. Data were plotted on GraphPad Prism 8. The spectra were noise reduced using the instrument algorithm and corrected for the buffer contributions without peak shape distortion and without altering peak maxima or intensities.

### Macrophage Culture

The murine macrophage cell line RAW 264.7 was maintained in RPMI-1640 medium (Gibco) supplemented with 10% heat inactivated fetal bovine serum at 37°C in a humidified atmosphere containing 5% CO_2_. The adherent cells were routinely passaged using a cell scrapper at around 80% confluence.

### In vitro protein translation in presence of wild type and mutant LmGAPDH

Total RNA was isolated from LPS (1μg/ml) treated RAW 264.7 cells for 6 hours using Trizol reagent (Invitrogen) following the manufacturer’s protocol. From total RNA, mRNA was segregated using PolyATtract® mRNA Isolation System (Promega) in accordance with the manufacturer’s protocol. The mRNA was incubated with wild type and mutant GAPDH proteins separately for 30 mins, then the mixture was subjected to *in vitro* translation by addition of 35 μl of rabbit reticulocyte lysate, nuclease treated (Promega), 1μl of RNasin® ribonuclease inhibitor (Promega), 1μl of 1 mM complete amino acid mixture (Promega), 1μl of Transcend^™^ tRNA (precharged, biotin-labeled-lysine tRNA) (Promega) in a 50μl reaction for 90 mins at 30°C. A 48μl aliquot was subjected to immunoprecipitation with TNF-α monoclonal antibody (Abcam) and 30 µl of Pierce^™^ protein A/G magnetic beads (Thermo Scientific). IP buffer was composed of 20 mM Tris-HCl (pH-7.5), 150 mM KCl, 5 mM MgCl_2_, 1 mM DTT and 1X PMSF.

Immunoprecipitated proteins as well as the 2 µl aliquot which was not subjected to immunoprecipitation, were separately resolved on 10% SDS-PAGE. The gel was analyzed with Transcend^™^ Non-Radioactive Translation Detection System having Streptavidin-Alkaline Phosphatase (Promega)

### Statistical analysis

All data were expressed as the mean ± SD from at least three independent experiments. Statistical analysis for parametric data was calculated by Student’s t test or analysis of variance (ANOVA) wherever applicable using GraphPad Prism 8.0 software. A p value of less than 0.05 was considered statistically significant.

## Data Availability Statement

All data generated and analyzed during this study are contained within the article.

## Supplementary data

This article contains supplementary data.

## Acknowledgments

This work was supported by Council of Scientific and Industrial Research (CSIR) NCP Project MLP-136 and Department of Science and Technology (CRG/2021/000421), CSIR fellowships (to G.C.) and University Grants Commission Fellowships (P.P., S.D. and Y.D.).

Conflict of interest: The authors declare that they have no conflicts of interest with the content of this article.

## Author contributions

Conceived and designed the experiments: S.A. Performed the experiments: P.P., S.D., Y. D., G. C. and S.A. Analyzed the data: P.P. and S.A. Contributed reagents/materials/analysis tools: S.A. Wrote the paper: S.A.

## List of Abbreviations

LmGAPDH: Glyceraldehyde-3-phosphate dehydrogenase from *Leishmania major*
WT: wild type enzyme
TNF-α: Tumor necrosis factor alpha
EV: extracellular vesicle
TNF-α ARE: AU-rich elements in 3’-untranslated regions of TNF-α mRNA
mwPIFE: protein induced fluorescence enhancement in a microwell plate
REMSA: RNA Electrophoretic mobility shift assay.

