## Supplemental figure and table for "Mapping of residues in leishmanial glyceraldehyde-3-phosphate dehydrogenase crucial for binding with 3’-UTR of TNF-α mRNA"

**Supplementary data**

**Table S1. Oligonucleotides used in this manuscript.**

| Primer | Primer sequence (underline indicate mutation sites) |
| --- | --- |
| R13A Forward | AACGGCTTCGGCGCCATTGGTCGCATG |
| R13A Reverse | CATGCGACCAATGGCGCCGAAGCCGTT |
| R13K Forward | TCAACGGCTTCGGCAAGATTGGTCGCATGGT |
| R13K Reverse | ACCATGCGACCAATCTTGCCGAAGCCGTTGA |
| I14A Forward | GGCTTCGGCCGCGCTGGTCGCATGGTG |
| I14A Reverse | CACCATGCGACCAGCGCGGCCGAAGCC |
| R16A Forward | GGCCGCATTGGTGCCATGGTGCTTCAG |
| R16A Reverse | CTGAAGCACCATGGCACCAATGCGGCC |
| D39A Forward | GTCGTCGCTGTCGTGGCCATGAGCACGAATGCTGAA |
| D39A Reverse | TTCAGCATTCGTGCTCATGGCCACGACAGCGACGAC |
| T112A Forward | TACGTGATCGAGTCTGCTGGCCTGTTCACGGACAAG |
| T112A Reverse | CTTGTCCGTGAACAGGCCAGCAGACTCGATCACGTA |
| N336A Forward | GTCTCGTGGTATGACGCCGAGTGGGGCTACTCGCAC |
| N336A Reverse | GTGCGAGTAGCCCCACTCGGCGTCATACCACGAGAC |

**Supplementary data**

**Figure S1. % Shifted band intensity of mutant protein-RNA complexes versus the concentration of mutant LmGAPDH.** Mutant LmGAPDH dependent shifted band Intensity of Fig. 6. Band intensity was quantified by ImageJ software (NIH). Error bars represent the SD from three independent experiments. Shifted band Intensity of I14A is not concentration dependent. Calculated K_D_ values for each mutant are: D39A (21 ± 1.8 μM), R16A (4.6 ± 1.1 μM) and T112A (8.6 ± 1.2 μM).

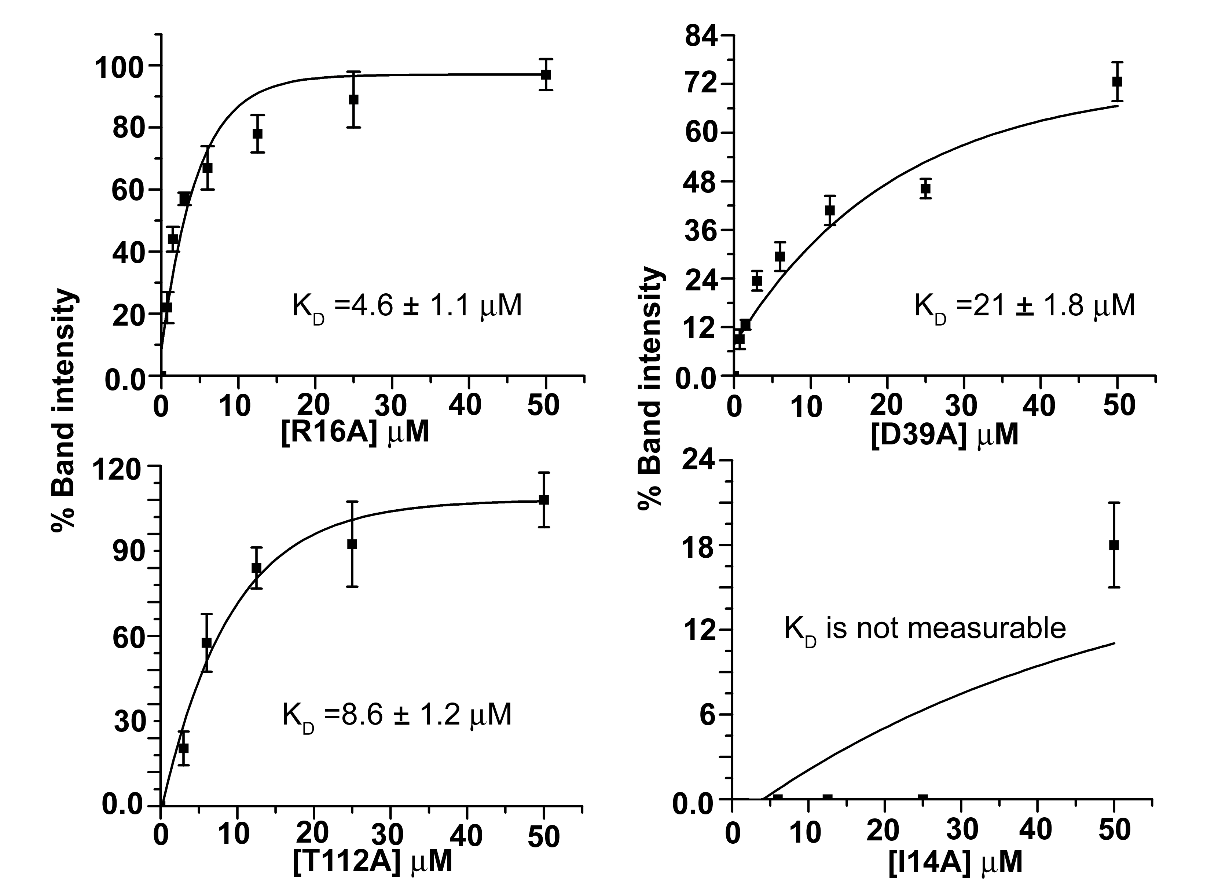
